# Kinetic modeling of ^18^F-PI-2620 binding in the brain using an image-derived input function with total-body PET

**DOI:** 10.1101/2024.07.02.601764

**Authors:** Anjan Bhattarai, Emily Nicole Holy, Yiran Wang, Benjamin A. Spencer, Guobao Wang, Charles DeCarli, Audrey P. Fan

**Author notes:** Corresponding author information: Correspondence to: Anjan Bhattarai; Department of Neurology and Department of Biomedical Engineering, University of California Davis, 1590 Drew Avenue, Unit #100 Davis, CA 95618, USA.

## Abstract

Accurate quantification of tau binding from ^18^F-PI-2620 PET requires kinetic modeling and an input function. Here, we implemented a non-invasive Image-derived input function (IDIF) derived using the state-of-the-art total-body uEXPLORER PET/CT scanner to quantify tau binding and tracer delivery rate from ^18^F-PI-2620 in the brain. Additionally, we explored the impact of scan duration on the quantification of kinetic parameters. Total-body PET dynamic data from 15 elderly participants were acquired. Time-activity curves from the grey matter regions of interest (ROIs) were fitted to the two-tissue compartmental model (2TCM) using a subject-specific IDIF derived from the descending aorta. ROI-specific kinetic parameters were estimated for different scan durations ranging from 10 to 90 minutes. Logan graphical analysis was also used to estimate the total distribution volume (V_T_). Differences in kinetic parameters were observed between ROIs, including significant reduction in tracer delivery rate (K_1_) in the medial temporal lobe. All kinetic parameters remained relatively stable after the 60-minute scan window across all ROIs, with K_1_ showing high stability after 30 minutes of scan duration. Excellent correlation was observed between V_T_ estimated using 2TCM and Logan plot analysis. This study demonstrated the utility of IDIF with total-body PET in investigating ^18^F-PI-2620 kinetics in the brain.

## Introduction

^18^F-PI-2620 is a second-generation PET radiotracer that has high binding affinity for tau protein aggregates in the brain, a key pathological feature in Alzheimer disease (AD), progressive supranuclear palsy, and other taopathies ^1–4^. Tau overload in the brain has been suggested to play a role in synaptic degeneration and neuronal loss ^5^. Initial clinical investigations and visual assessments have shown increased ^18^F-PI-2620 binding in the brain in individuals with AD ^4,6^. However, there is a need for studies investigating subtle change in tau binding in early-stage AD and elderly individuals at risk of AD. Accurate, non-invasive quantification of ^18^F-PI2620 tau load in the brain is critical in research as well as clinical trials with therapeutic interventions.

Conventionally, standard uptake value ratio (SUVR) has been used to evaluate tracer distribution in the brain ^6–8^. SUVR is a semiquantitative measure and is normalized to tracer uptake in a reference region, e.g. the inferior cerebellum for tau PET. SUVR comparisons across disease groups assume that tracer uptake in a reference region remains unchanged despite the presence of pathologies. SUVR does not take into account other pharmacokinetic components, such as blood volume, tracer metabolism, and blood flow, which may influence the overall uptake in the targeted region. Furthermore, SUVR does not characterize the transport information of the tracer.

Accurate quantification of tau binding from ^18^F-PI-2620 PET requires kinetic modeling and an input function. Conventionally, the arterial input function (AIF), which requires arterial blood sampling, has been used for kinetic modeling ^9^. Although AIF is considered the gold-standard approach for kinetic parameter estimation, it is an invasive approach and may cause discomfort and discourage participation in studies. AIF is also susceptible to sampling errors, delay, and dispersion, which may often lead to inaccurate estimation of kinetic parameters. Non-invasive image-derived input function (IDIF), derived using dynamic PET imaging, has emerged as an alternative to AIF ^9–11^. IDIF eliminates the need for arterial cannulation, manual blood handling, and analyses, thereby minimizing patient discomfort and reducing the exposure of additional radiation to the research personnel involved in data acquisition ^11^. It is generally challenging to obtain a reliable IDIF for dynamic brain PET imaging with a conventional short scanner due the limited axial field-of-view that does not cover a major blood pool ^12^.

A recent development in PET imaging, the total-body uEXPLORER PET/CT scanner, offers an opportunity to generate total-body dynamic images with its long axial field-of-view, high photon collection efficiency, and high spatial resolution ^13,14^. More importantly, it enables us to extract IDIF from large blood pools such as the aorta and ventricles, which are less susceptible to partial volume effects compared to the conventionally suggested carotid artery. Our previous amyloid study with total-body PET demonstrated the utility of descending aorta IDIF for quantifying grey matter kinetics in the brain ^15^.

Kinetic modeling with ^18^F-PI-2620 requires dynamic PET data with relatively longer scan time (generally over one hour). However, such long scanning durations are cost-intensive and can be challenging for elderly participants. A previous study suggested that a shorter scan time (i.e., 40 minutes) can be sufficient for the measurements of semi quantitative SUVR in the brain in individuals with progressive supranuclear palsy (PSP) ^16^. There is a need for studies investigating the sensitivity and efficacy of shorter scan durations for estimating ^18^F-PI-2620 grey matter kinetics, quantified through a compartmental modeling approach, particularly in individuals with AD or those at risk of developing it.

Earlier findings have identified regional differences in ^18^F-PI-2620 uptake in AD cohort ^4,6^. The medial temporal lobe, including the hippocampus, entorhinal cortex, and amygdala, has been reported to have higher tau accumulation in AD ^6^. Tau kinetics in the grey matter regions known to be impacted based on Braak staging ^17^, in elderly cohorts at risk of AD, warrant further investigation. Investigating early disease stage tau kinetics with specific pharmacokinetic components is critical in research, as well as for clinical trials aimed at therapeutic intervention to delay disease progression.

This study aimed to implement the IDIF derived from a major blood pool using the total-body uEXPLORER PET/CT scanner to quantify ^18^F-PI-2620 kinetics with kinetic modeling of key brain grey matter regions in elderly individuals. Additionally, we investigated how scan duration influences the estimation of micro and macro kinetic parameters across grey matter region of interest (ROIs), quantified through two-tissue compartment modeling (2TCM). Furthermore, total distribution volume (V_T_), estimated using 2TCM was compared to the simpler graphical Logan plot analysis, both using IDIF. Finally, the study aimed to explore differences in the kinetic parameters across ROIs.

## Materials and methods

### Participants

Ethics approval for this study was obtained from the UC Davis Institutional Review Board. All the participants recruited in the study were part of the UC Davis Alzheimer’s Disease Research Center, over 65 years of age and gave written informed consent. Eligibility criteria included the ability to undergo an MRI and a known cognitive status based on clinical assessment and neuropsychological testing. Individuals with pacemakers, brain tumors, alcoholism, and/or those who were not able to lie still for 90 minutes were not included in this study. The study cohort included 15 individuals (age = 66-92 years; male = 9; female = 6), including 11 Cognitively unimpaired individuals, 2 individuals with mild cognitive impairment, and 2 with Alzheimer’s disease. Our cohort included 2 amyloid positive participants but did not include tau positive individuals.

### Data acquisition

#### PET

Dynamic total-body ^18^F-PI-2620 PET images were acquired for 90 minutes using the uEXPLORER and reconstructed with ordered-subset expectation-maximization (OSEM), (resolution = 2.344 mm isotropic voxels, four iterations, 20 subsets, with all standard data corrections except resolution modelling, and no post-reconstruction smoothing, following the UC Davis clinical protocol ^18^. A high temporal resolution dynamic framing protocol was used: 30×2s, 12×10s, 7×60s, 16×300s. The average injected dose was 185.33 MBq.

#### MRI

MRI (T1-weighted) images for all the participants were acquired using a 3-Tesla Siemens Tim Trio scanner (Siemens, Erlangen, Germany) equipped with a 32-channel head coil. The T1-weighted magnetization prepared rapid gradient echo (MPRAGE) images were acquired using the following parameters: matrix size = 240×256, in-plane spatial resolution = 1mm, acquisition time = 9 min 14 s, repetition time = 2300ms, echo time = 2.98ms, flip angle = 9°, inversion time = 900-1100s, voxel size = 1 × 1 × 1 mm^3^, 176 sagittal slices with thickness=1.0-1.2 mm. PET and MRI data were acquired on average 2.2 ± 1.2 years apart from each other.

### Motion correction and registration

Individual total-body dynamic PET images were brain-cropped using PMOD (PMOD Technologies, LLC), and motion corrected across time frames (from frame 50, i.e., longer than 10 minutes of scan) using FSL’s *mcflirt*. The 4D motion-corrected dynamic PET images were linearly (affine, with the default 12 degrees of freedom) registered to their respective T1-weighted (T1W) brain images using FSL’s flirt algorithm ^19^.

### Grey matter ROIs

Grey mater region of interests (ROIs), which are known to be impacted in AD based on Braak staging ^6,17^, were obtained from the cortical parcellation and subcortical segmentation (*recon-all*) of anatomical T1-weighted image using FreeSurfer (v6.0.0) ^20^.We focused on three subject-specific bilateral ROIs from the resulting *aseg* and *aparc* atlases ^21,22^. The ROIs were medial temporal lobe (combination of entorhinal, hippocampus, and amygdala), posterior cingulate (combination of isthmus cingulate and posterior cingulate), and lateral parietal cortex (combination of inferior parietal, superior parietal, and supramarginal) ^6^.

Bilateral grey matter cerebellum ROIs were obtained using FreeSurfer and manually edited to remove the superior portion (using the primary fissure as the posterior boundary), given known off-target binding issues with the superior cerebellum ^6^. The inferior cerebellum was used as a reference region for SUVR and distribution volume ratio (DVR) estimation. SUVR for 60-90 minutes was calculated as the ratio of activity in the target ROI to the mean activity in the reference region ROI (inferior cerebellum).

FreeSurfer was used to estimate the total (bilateral) hippocampal volume for each participant. Hippocampal volume was normalized by total intracranial volume, and its logarithm was estimated.

### Image derived input function

Individual image derived input functions (IDIFs) were extracted from the descending aorta using total-body dynamic PET images **(Figure 1).** The descending aorta ROIs were manually drawn using FSL image viewer, following a similar approach as employed in our previous studies ^15,23^.

**Figure 1:**
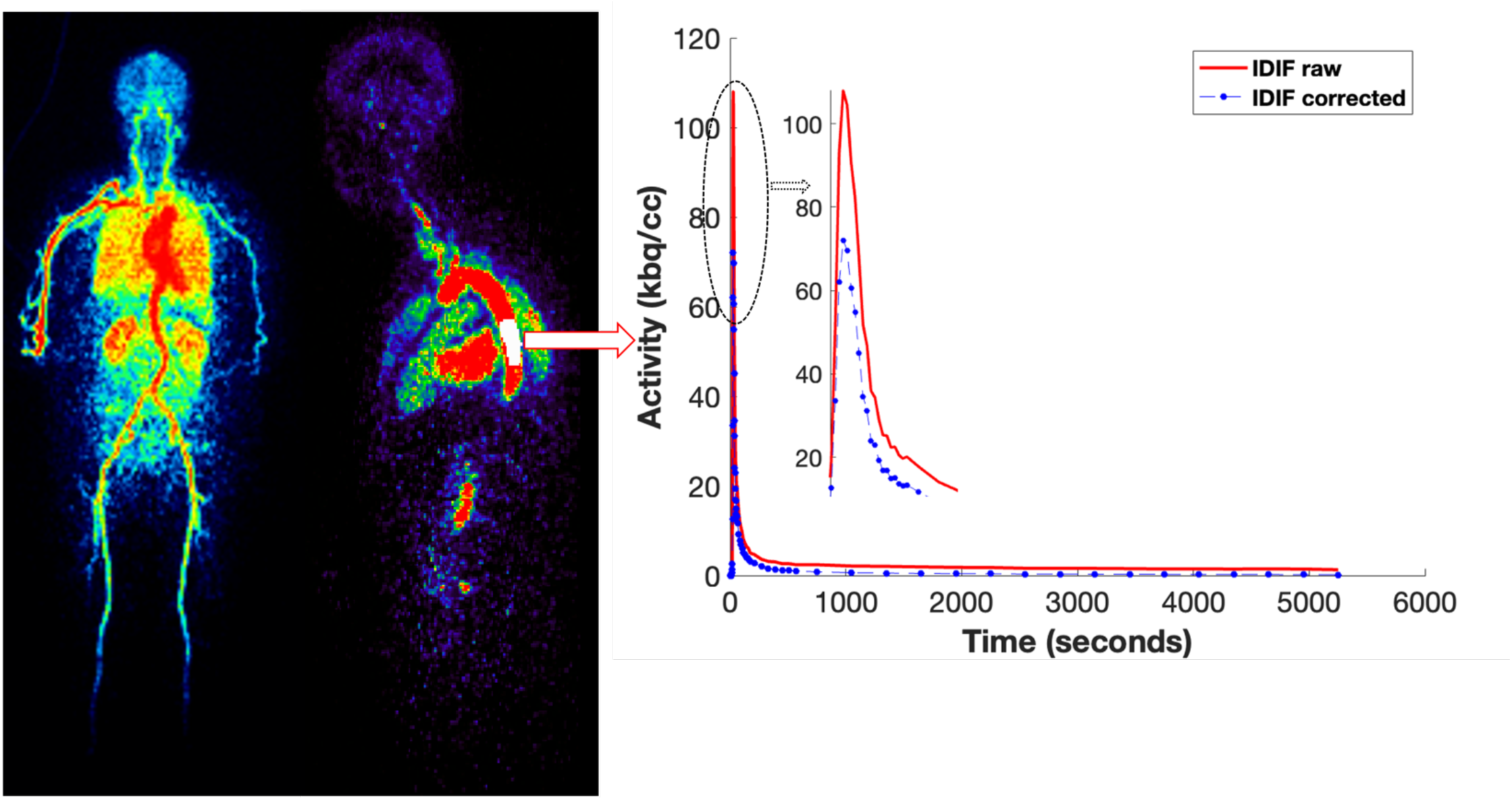
Input function derived from the descending aorta. IDIF: Image-derived input function; IDIF corrected: Plasma and metabolite corrected IDIF using population-based fractions over time. IDIF peak is zoomed for visualization.

These manual ROIs were eroded by 1mm in all dimensions to avoid the inclusion of vessel walls. The descending aorta time-activity curves were extracted and corrected for plasma and metabolite fractions prior to kinetic modeling. Plasma correction was performed using the total plasma to whole blood ratio data acquired from a previous ^18^F-PI-2620 study investigating tau deposits in the human brain^3^. A bi-exponential function describing the parent fraction (Equation 1), i.e. unmetabolized ^18^F-PI-2620 over time, was applied for metabolite correction ^4^.

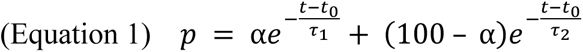

where *p* is percentage parent fraction; *t* is time after injection, in minutes; and α, τ_1_, τ_2_, and *t*_0_ are the model parameters (**Supplement Figure 2**).

### Compartmental modeling

Dynamic time-activity curves (TACs) for each brain ROI were fitted to a reversible two-tissue compartmental model (2TCM) **(Figure 2)**, using a subject-specific IDIF derived from the descending aorta. A joint estimation-based approach was used to correct for delay of tracer arrival from the input function to the tissues of interest ^24^.

**Figure 2:**
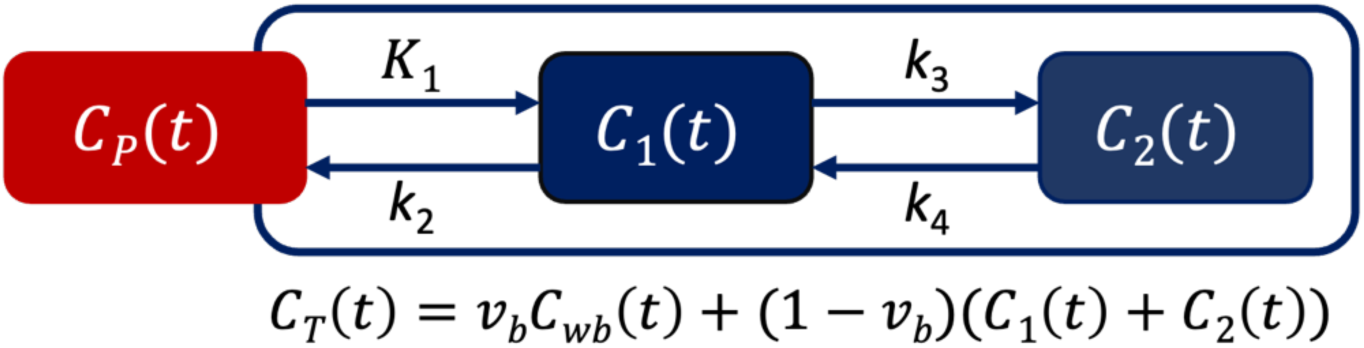
A reversible two-tissue compartmental model for ^18^F-PI-2620 kinetic modeling in the brain. *C_p_(t)* represents the activity of ^18^F-PI-2620 in the blood plasma. *K*_1_, *k*_2_, *k*_3_, and *k*_4_ are rate constants representing the transfer of the tracer between plasma and tissue compartments, as well as between tissue compartments. *C_1_(t)* and *C_2_(t)* represent the concentration of the tracer in brain tissue compartments. *C_T_(t)* represents the total ^18^F-PI-2620 activity measured by PET in the brain compartments. *v_b_* is fractional blood volume, and *C_wb_(t)* is the tracer activity in whole blood.

The following kinetic parameters were estimated for each ROI: blood fraction volume (v_b_ [(ml/ml)]), rate constants (K_1_ [mL/cm^3^/min]), k_2_ [min^−1^], k_3_ [min^−1^], k_4_ [min^−1^]), macro kinetic parameters (i.e., V_T_ [ml/cm^3^], and distribution volume ratio DVR [unitless]), and tracer arrival delay (sec) ^24,25^. The total distribution volume V_T_ was estimated as: 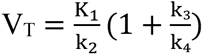. The DVR using the inferior cerebellum as a reference region was computed as V_T_/ V_T(ref)_, in which V_T_ represents the total distribution volume in the target ROI and V_T(ref)_ represents the total distribution volume in the reference region ^3^. Additionally, SUVR corresponding to 60-90 minutes post injection was calculated using the inferior cerebellum as a reference region.

### Logan Plot analysis

Logan graphical analysis (t*=20 min) was used to estimate total distribution volume (V_T_), using the image-derived input function *C*_P_(t) and the brain time-activity curve, *C*_T_(t), for each ROI (Equation 2) ^26^.

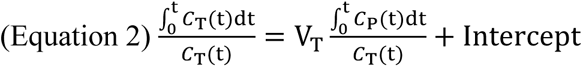

### Scan duration

Kinetic parameters were estimated for different scan durations (from the start of injection) ranging from 10 to 90 minutes, in intervals of 10 minutes. Quantification at all ROIs were performed using dynamic data acquired at different durations and was compared with the results from the full 90-minute scan.

### Statistical analysis

Kinetic parameters (mean ± standard deviation) for all ROIs, obtained with the full 90-minute data and shorter scan durations, were calculated. Kinetic measures estimated with shorter scan durations were correlated (using Pearson’s correlation across all participants) with the respective measures estimated from the full 90-minute dynamic scan.

A linear mixed-effects model (using MATLAB’s *fitlme* function) was employed to investigate the relationship between kinetic measures (quantified with 90-minute data) and ROIs, accounting for subject variances. False Discovery Rate (FDR) was applied for multiple comparisons correction using the Benjamini-Hochberg procedure (using MATLAB’s *mafdr* function). Significance was determined based on the FDR-adjusted *q-value* threshold of *q* < 0.05. Additionally, as an exploratory analysis, participants’ cognitive status was incorporated as a covariate in the mixed-effects model.

Associations between V_T_ estimated using 2TCM and Logan plot analysis for each ROI were estimated using Pearson’s correlation. The significance (*p-value* at the significance level *α* = 0.05) of the association using simple linear regression (using MATLAB’s *polyfit* function) was reported. Associations between DVR and SUVR measures were estimated using Pearson’s correlation. Bilateral hippocampal volumes (logarithm of normalized volume) from MRI segmentations were also correlated with K_1_ measures of the medial temporal cortex. Additionally, a linear mixed-effects model was used to investigate the relationship between SUVR and ROIs, accounting for subject variances.

## Results

### Compartmental modeling

Two-tissue compartmental modeling with IDIF, corrected for plasma and metabolites, and accounting for time delay through joint estimation, demonstrated high-quality fits of ^18^F-PI-2620 binding across grey matter ROIs, for all individuals. **Figure 3** displays time-activity curves for all three ROIs of an 81-year-old participant with AD, revealing strong model fits to our data.

**Figure 3:**
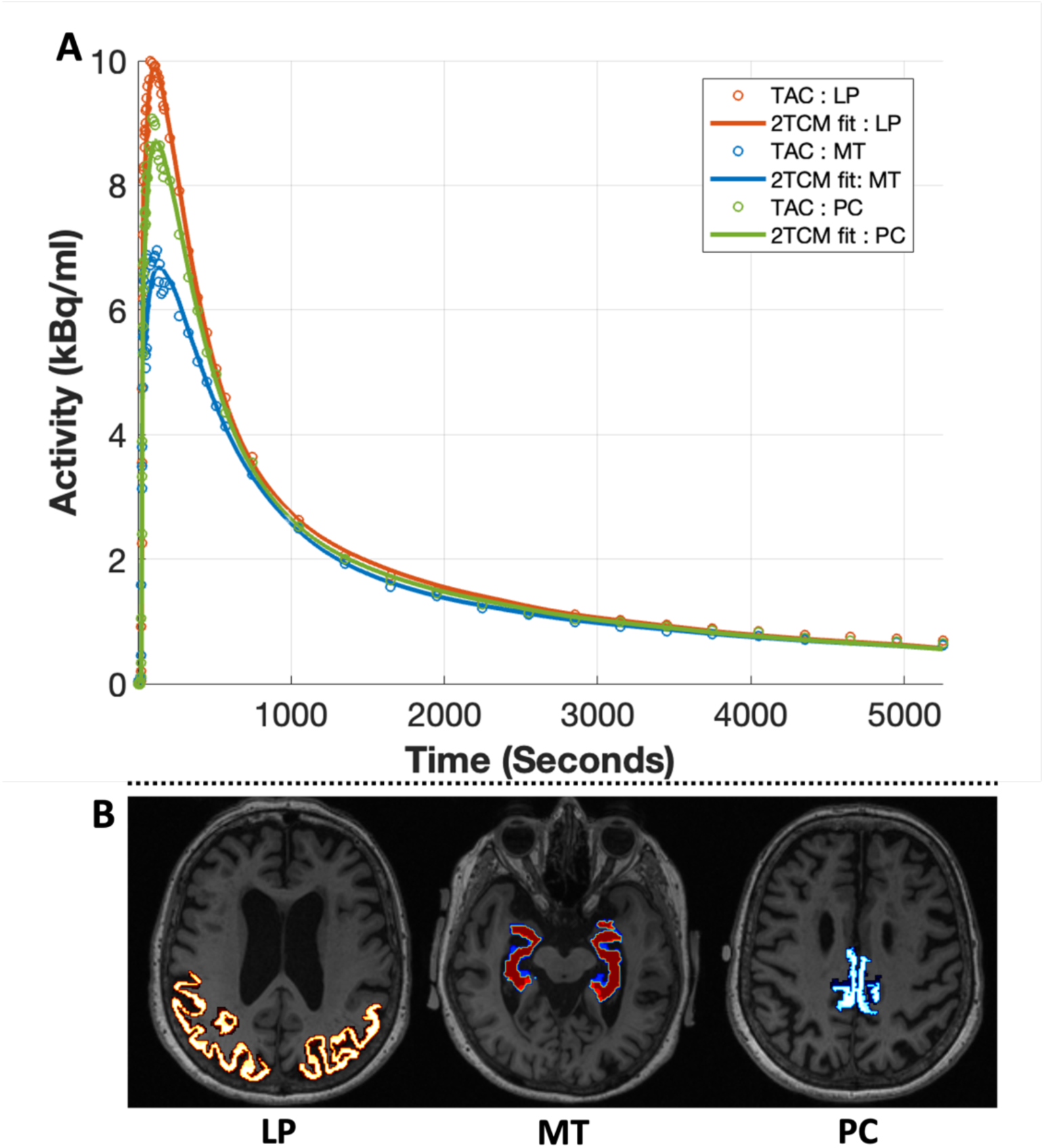
Two-tissue compartmental modeling (2TCM) curve fitting. **A**: Curve fitting for grey matter ROIs, known to be vulnerable in AD and dementia, in an individual (81-year-old female, amyloid-positive, tau-negative, diagnosed with AD). **B**: Grey matter ROIs overlaid on T1W image. TAC: Time-activity curve; LP: lateral parietal; MT: Medial Temporal; PC: Posterior cingulate.

### Regional differences in kinetic parameters

Differences in quantified kinetic parameters were observed across ROIs (**Table 1**). The linear mixed-effect model analyses revealed significant differences in v_b_ between LP and MT (*β*= 0.010, *q*<0.001), and between LP and PC *(β* = 0.008, *q* < 0.001). No significant differences in V_T_ were observed across ROIs.

**Table 1:** Quantified Kinetic parameters across all 15 participants in the three key ROIs. V_T_Logan is V_T_ estimated with graphical logan plot analysis. *Significant differences at FDR *q*<0.05 according to linear mixed-effects model analysis, comparing the measures across ROIs: *V*_b_: LP vs. MT, LP vs. PC; *K*_1_: LP vs. MT, MT vs. PC; *k*_2_: LP vs. MT, MT vs. PC; *Delay*: LP vs. MT, MT vs. PC. See supplementary material for full details of the results.

| ROIs | Kinetic parameters (mean± std) |  |  |  |  |  |  |  |  |
| --- | --- | --- | --- | --- | --- | --- | --- | --- | --- |
| | $v_b$ (ml/ml) | $K_1$(mL.cm<sup>-3</sup>.min<sup>-1</sup>) | $k_2$ (min<sup>-1</sup>) | $k_3$ (min<sup>-1</sup>) | $k_4$ (min<sup>-1</sup>) | Delay (s) | $V_T$ (mL.cm<sup>-3</sup>) | DVR | $V_{TLogan}$ (mL.cm<sup>-3</sup>) |
| LP | 0.029 ± 0.004* | 0.372 ± 0.043 | 0.245 ± 0.034 | 0.031 ± 0.019 | 0.083 ± 0.055 | 4.625 ± 0.514 | 2.111 ± 0.275 | 1.035 ± 0.075 | 2.179 ± 0.266 |
| MT | 0.038 ± 0.007* | 0.271 ± 0.036* | 0.190 ± 0.043* | 0.039 ± 0.047 | 0.070 ± 0.058 | 3.766 ± 0.902* | 2.159 ± 0.331 | 1.057 ± 0.106 | 2.216 ± 0.291 |
| PC | 0.037 ± 0.007 | 0.369 ± 0.042 | 0.253 ± 0.031 | 0.031 ± 0.017 | 0.076 ± 0.038 | 4.323 ± 0.625 | 2.081 ± 0.236 | 1.022 ± 0.075 | 2.168 ± 0.225 |

Notably, differences in the tracer delivery parameter K_1_ were observed across ROIs. K_1_ was significantly reduced in MT relative to both LP and PC (LP and MT: *β*= -0.100, *q* < 0.001; MT and PC: *β*= 0.098, *q* < 0.001). No significant differences in K_1_ were observed between LP and PC.

Similarly, significantly reduced k_2_ was observed in MT compared to LP and PC **(Table 1).** Furthermore, significantly reduced tracer arrival time delay was observed in MT compared to LP (*β*= -0.859, *q* < 0.001). The observed regional differences did not change when participants’ cognitive status was incorporated as a covariate in the mixed-effects model. No group differences were observed between cognitively impaired and unimpaired groups. Excellent correlation was observed between V_T_ estimated using 2TCM and Logan plot analysis, across all ROIs (LP: r = 0.99, *p*<0.001; MT: r = 0.96, *p*<0.001; PC: r = 0.99, *p* < 0.001) (**Figure 4**).

**Figure 4:**
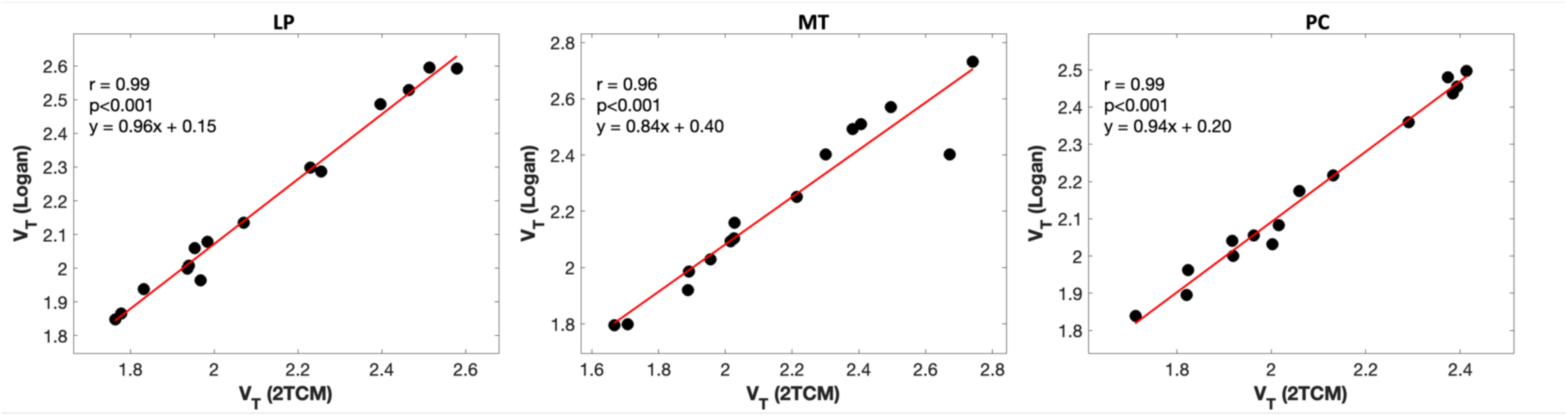
Comparison of V_T_ estimation between 2TCM and Logan analyses across ROIs. LP: Lateral parietal; MT: Medial temporal; PC: Posterior cingulate.

### SUVR and its association with DVR

Significantly increased SUVR was observed in MT compared to LP and PC (LP and MT: *β*= 0.120, *p<0.001)*; MT and PC: *β*= -0.110, *p* < 0.001). Significant associations were observed between DVR and SUVR measures across all ROIs. (LP: r = 0.60, *p*= 0.02; MT: r = 0.82, *p*<0.001; PC: r = 0.63, *p* =0.01). (**Supplement Figure 1)**.

### Effect of scan duration on kinetic parameters

All kinetic parameters remained relatively stable after the 60-minute scan window across all ROIs, demonstrating the significant correlation (r≥0.89; p<0.001) with the parameters derived from the full 90-minute data (**Figure 5 and 6**). K_1_ was highly stable (r≥0.92; p<0.001) after 30 minutes of scan duration across all ROIs.

**Figure 5:**
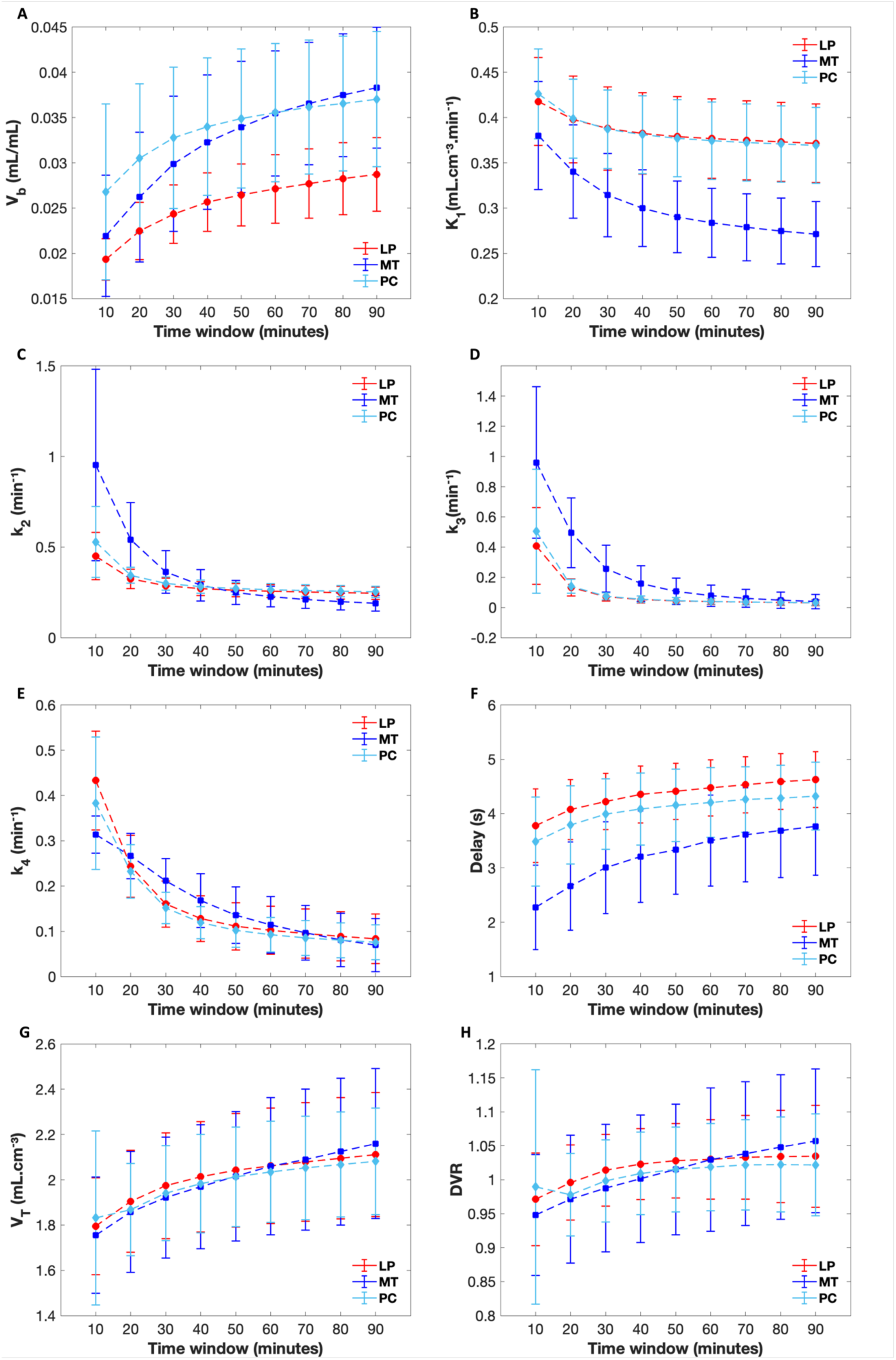
Kinetic parameters (K_1_, k_2_, k_3_, k_4_, delay, V_T_ and DVR) estimated for different scan durations. Error bars and means (connected with lines) are shown for each ROI, at each scan time. LP: Lateral Parietal; MT: Medial temporal; PC: Posterior cingulate.

**Figure 6:**
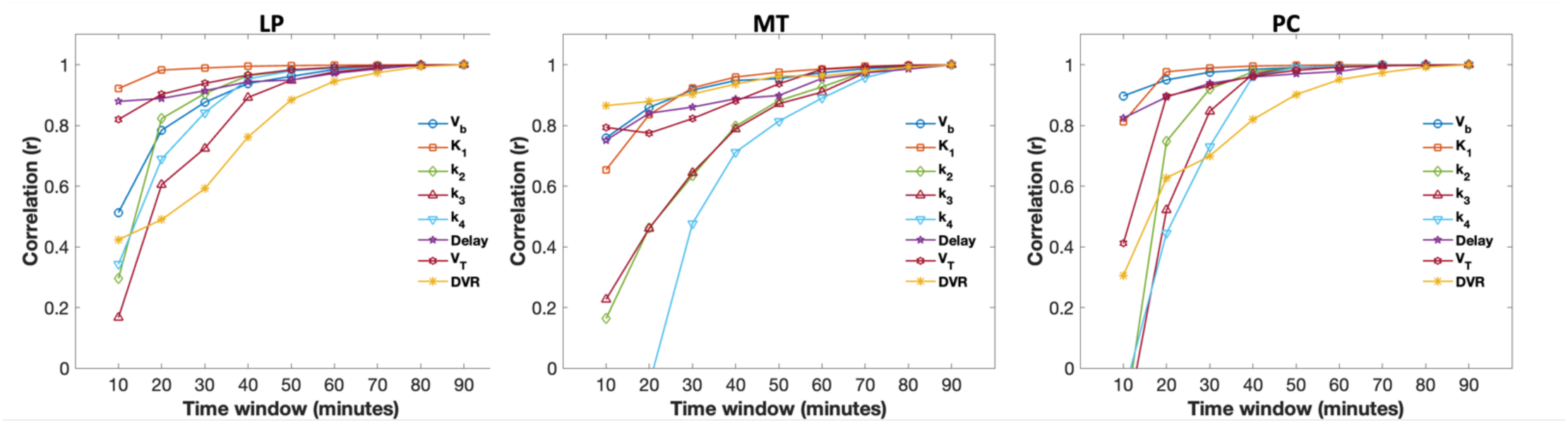
Evolution of correlations over scan time. Correlation (r) is the Pearson’s correlation between kinetic parameters estimated at each shorter time window and full 90-minute scan.

For PC, strong agreement (r≥0.90; p<0.001) was observed between the kinetic parameters estimated using scan durations of 50 minutes or longer and those derived from the full 90-minute data (**Figure 6**). Similarly, for LP, strong agreements with the 90-minute data (r≥0.88; p<0.001) were observed after 50 minutes of scanning. For MT, compared to the other two ROIs, the parameters exhibited relatively more instability (**Figure 5**). However, strong correlation (r≥0.89; p<0.001) with the full 90-minute dataset was observed after 60 minutes of scanning. It is noted that V_T_ and DVR (especially in MT) compared to other parameters exhibited more instability (as observed in the evolution curves) even after 60 minutes of scanning (**Figure 5, G, H**).

### Associations between kinetic parameters

Our exploratory analyses (with linear mixed-effects model) revealed a significant association between perfusion K_1_ (*β* = 0.03, *p* = 0.012) and tracer arrival delay, including the measures computed across all ROIs. No significant associations between K_1_ and V_T_ (*β* = -0.16, *p* = 0.702) were observed. No significant correlation was observed between hippocampal volume and the K_1_ values (r = -0.37, *p* = 0.17) in MT.

## Discussion

This study implemented a non-invasive descending aorta IDIF, derived using a state-of-the-art total-body uEXPLORER PET/CT scanner, to quantify tau binding and tracer delivery rate from ^18^F-PI-2620 across grey matter ROIs in elderly individuals. Regional differences in tracer delivery rate were observed, suggesting medial temporal lobe to be susceptible to change in blood flow dynamics. We investigated the impact of scan duration on estimating kinetic parameters, suggesting a 60-minute scan may be required for reliable estimation of pharmacokinetic parameters (i.e., based on comparison with kinetic parameters obtained with a 90-minute scan), while a 30-minute scan may be sufficient for quantifying perfusion. A reversible 2TCM approach was used to quantify multiple micro and macro parameters, thus providing more insights into physiology. At the same time, the findings suggest that the graphical Logan plot analysis may be sufficient for estimating the macro kinetic parameter, total distribution volume, though it cannot estimate K_1_. Together, these findings highlight the utility of IDIF with total-body PET in clinical research.

This study focused on investigating tau deposition in an elderly cohort at risk of dementia. Earlier findings suggest that early tau deposition in cognitively unimpaired individuals may exhibit a divergent cortical tau PET pattern ^27^ and that tau PET signal in medial temporal lobe regions correlates with poorer cognitive scores in memory tests in unimpaired individuals ^28^. Investigating early-stage tau PET deposition and kinetics is critical for clinical research, as well as for selecting participants for clinical trials aimed at preventing or halting disease progression. Our findings in tau-negative individuals could also potentially serve as a quantitative reference for future ^18^F-PI-2620 kinetic modeling studies.

Conventionally used SUVR measures with static PET do not provide absolute quantification of tracer distribution in the targeted tissue. In this study, we observed a moderate correlation between SUVR and DVR measures in the lateral parietal and posterior cingulate ROIs. While SUVR is a commonly used metric, it does not account for dynamic changes in tracer distribution over time and relies on a reference region, which may contribute inaccuracies. Kinetic modeling allows for absolute and accurate quantification of micro and macro kinetic parameters without the need for a reference region. Kinetic modeling requires an input function ^29^. Although the arterial input function is still considered the gold standard for kinetic modeling, it requires blood sampling and is invasive in nature. Blood arterial inputs are also susceptible to sampling errors, delay, and dispersion ^10,30^.

IDIF extraction is an alternative approach but can be very challenging when small-diameter, convoluted vessels, and complex surrounding structures are present inside the field-of-view. A recent ^18^F-PI-2620 brain study explored the utility of IDIF against AIF for estimating a macroparameter, V_T_, with the graphical Logan plot analysis ^30^. However, the study utilized the carotid artery as an IDIF for the brain, which has a diameter of approximately 4 to 5 mm and is smaller than the spatial resolution of most of the clinical PET scanners (∼ 5 mm) ^10^. The limited spatial resolution may introduce partial volume effects in the IDIF and may lead to incorrect parameter estimation ^31^. We focused on a more comprehensive and accurate kinetic compartmental modeling approach to examine tracer distribution, which accounts for several pharmacokinetic measures (e.g., K_1_) in addition to V_T_. More importantly, this approach utilized a larger blood pool as an IDIF, correcting for plasma and metabolites, for an accurate estimation of kinetic parameters.

The total-body uEXPLORER PET/CT scanner offers the opportunity to extract IDIF from larger blood pools including ventricles and aorta, which would not be possible using the conventional scanners with shorter FOV. We used the descending aorta as the IDIF, which is relatively less susceptible to partial volume effects. Anatomically, the descending aorta is closer to the posterior chest wall, which protects it from cardiac and respiratory motion, making it an optimal choice for IDIF in kinetic modeling studies ^32^. Furthermore, the descending aorta, instead of the ascending aorta, was selected as the IDIF in this study to avoid additional complexity and potential errors (associated with visibility of the ascending aorta in dynamic PET data) while manually extracting the ROIs. We have previously demonstrated the utility of the descending aorta IDIF in our amyloid PET study ^15^.

This study also benefitted from the excellent spatial resolution (voxel size of 2.34 mm^3^) and high temporal resolution framing (2 second initial frames) of the total-body uEXPLORER PET/CT scanner. The enhanced temporal resolution enabled accurate estimation of tracer distribution in early frames. Earlier studies using ^18^F-PI-2620 had significantly lower temporal resolution, typically around 30 seconds ^3,4^.

In our study, two-tissue compartmental modeling with the descending aorta IDIF demonstrated high-quality fitting of ^18^F-PI-2620 binding across grey matter ROIs. Regional differences in kinetic micro and macro parameters were observed, but no differences in total distribution volume (V_T_) were observed. Notably, reduced tracer delivery rate (K_1_) was observed in MT, which may indicate reduced perfusion in that region. Our results corroborate earlier findings that have associated hypoperfusion in the medial temporal lobes with cognitive decline, characterized by decreased performance on memory tests, and increased risk for AD ^33,34^. We observed reduced tracer arrival time (i.e., delay) and increased blood fraction volume in MT. We speculate that these regional differences in tracer delivery and blood flow dynamics may reflect physiological and metabolic differences in MT, which is known to be vulnerable in aging and dementia. Significant associations were also found between K_1_ and arrival delay, suggesting that potential differences in cerebral blood flow and grey matter vascular dynamics could impact both K_1_ and tracer transport. Although we observed a strong correlation between the macroparameter V_T_ as estimated by Logan graphical approach and 2TCM, the 2TCM compartment modeling in this study allowed us to estimate several microparameters, providing more insights into tracer kinetics.

These findings warrant validation in a larger cohort with additional cognitive impaired and tau-positive individuals. Our primary goal was to implement aorta-derived IDIF in kinetic modeling; however, there is an avenue for future research to apply our approach to investigate differences in grey matter tracer dynamics between tau-positive and tau-negative individuals. Although not statistically significant, we also observed a trend of increasing K_1_ with hippocampal volume in MT. Future research with longitudinal quantitative tau PET follow-up could investigate whether structural changes in the hippocampus may impact cerebral perfusion and tau levels in healthy aging and early AD.

The K_1_ parameter appeared to be highly stable (compared with K_1_ estimated from the full 90 minutes scan) after 30 minutes of scan duration across all ROIs, suggesting that shorter time durations may be sufficient for the quantification of perfusion measures across grey matter ROIs. All other kinetic parameters remained relatively stable after the 60-minute scan window across all ROIs. It is important to note that kinetic parameters in MT, especially V_T_ and DVR were less stable compared to other ROIs. These regional variations and parameters selection need to be accounted for when selecting the scanning protocols for ^18^F-PI-2620 kinetic modeling studies. However, our findings need to be validated in a cohort which includes tau-positive individuals. An earlier study suggested that a shorter scan time (i.e., 40 minutes) can be sufficient for the measurements of semi-quantitative SUVR in the brain in individuals with progressive supranuclear palsy (PSP). The impact of scan duration on the estimation of quantitative kinetic microparameters has not been previously studied for this tracer. Our findings suggest that 60-minute time window may be adequate for quantifying micro and macro kinetic parameters with two-tissue compartmental modeling in elderly cohort at risk of AD.

There are some limitations in our study. Our cohort did not include tau-positive individuals, and it was relatively small. IDIF for kinetic modeling is emerging and has the potential to replace the conventional invasive arterial blood sampling approach in a long term, however, the utility of IDIF for ^18^F-PI-2620 kinetic modeling requires further investigation and validation against blood sampling. We utilized the descending aorta IDIF in this study. We recommend that future studies explore the impact of IDIF selection (i.e., from other anatomical locations) in ^18^F-PI-2620 kinetic modeling studies. We did not consider the potential intersubject variability in ^18^F-PI-2620 metabolism for metabolite correction. ^18^F-PI-2620 metabolism may vary with the age or cognitive status, as slightly faster metabolism has been reported in AD compared to healthy subjects ^3^. This study implemented a bi-exponential function, as discussed in a previous study ^4^, to estimate the parent fraction for metabolite correction. A full examination of potential individual differences in ^18^F-PI-2620 metabolism, especially among aging, mild cognitive impairment, Alzheimer’s disease (AD), and control individuals, would facilitate a more accurate quantification and interpretation of tracer dynamics in the brain.

## Conclusion

This study implemented a descending aorta IDIF, derived using a novel total-body uEXPLORER PET/CT scanner, for the non-invasive quantification of ^18^F-PI-2620 in the brain of elderly individuals at risk of AD. The study used a reversible 2TCM and a graphical Logan plot analysis to quantify tracer kinetics. No significant differences in regional total distribution volume were observed, but there was a significant reduction in the tracer delivery rate, K_1_, in MT. Our findings suggest that a 60-minute scan window may be required for the reliable quantification of kinetic parameters using IDIF, whereas a 30-minute scan time may be sufficient for the quantification of K_1_. These findings need to be validated in a larger tau-positive AD cohort. Overall, the study demonstrated the utility of the uEXPLORER PET/CT scanner and kinetic modeling with IDIF in clinical research.

## Acknowledgements

We would like to thank all participants, their families, and the clinicians involved in this study. Without their help, this study would not have been possible.

## Author contributions statement

**Anjan Bhattarai:** Conceptualization, Formal Analysis, Investigation, Methodology, Software, Validation, Visualization, Writing -original draft, Writing -review and editing. **Emily Nicole Holy:** Methodology, Software, Visualization, Writing -review and editing. **Yiran Wang:** Methodology, Software, Writing -review and editing. **Benjamin A. Spencer:** Methodology, Software, Writing -review and editing. **Guobao Wang:** Methodology, Software, Writing -review and editing. **Charles DeCalrli:** Funding Acquisition, Project administration, Resources, Investigation, Writing -review and editing. **Audrey P. Fan:** Funding Acquisition, Conceptualization, Project administration, Resources, Investigation, Supervision, Methodology, Validation, Writing -review and editing.

## Declaration of conflicting interests

The author(s) declared no potential conflicts of interest with respect to the research, authorship, and/or publication of this article.

## Funding

This study is supported by ADRC center grant P30 AG072972.

## Data availability statement

The data from this study are not publicly available due to the privacy issues of the participants. Data will be shared upon reasonable request for research only after ethical approval for the specific project.

## Supplements

**Supplement Table 1:** Relationship between kinetic measures (quantified with 90-minute data) and ROIs, accounting for subject variance, estimated using a linear mixed-effects model. Significance was determined based on the FDR-adjusted q-value threshold of q < 0.05. Standard error (SE). t-statistics (tStat). Degree of freedom (DF). Confidence interval (CI).

| Measure | Comparison | Estimate | SE | tStat | DF | qValue | Lower CI | Upper CI |
| --- | --- | --- | --- | --- | --- | --- | --- | --- |
| **v<sub>b</sub>** | LP and MT | 0.010 | 0.001 | 6.591 | 42 | <0.001* | 0.007 | 0.013 |
|  | LP and PC | 0.008 | 0.001 | 5.714 | 42 | <0.001* | 0.005 | 0.011 |
|  | MT and PC | -0.001 | 0.001 | -0.877 | 42 | 0.517 | -0.004 | 0.002 |
| **K<sub>1</sub>** | LP and MT | -0.100 | 0.009 | -11.316 | 42 | <0.001* | -0.118 | -0.082 |
|  | LP and PC | -0.002 | 0.009 | -0.259 | 42 | 0.831 | -0.020 | 0.016 |
|  | MT and PC | 0.098 | 0.009 | 11.057 | 42 | <0.001* | 0.080 | 0.116 |
| **k<sub>2</sub>** | LP and MT | -0.055 | 0.008 | -6.799 | 42 | <0.001* | -0.071 | -0.039 |
|  | LP and PC | 0.008 | 0.008 | 0.968 | 42 | <0.001* | -0.008 | 0.024 |
|  | MT and PC | 0.063 | 0.008 | 7.768 | 42 | <0.001* | 0.047 | 0.079 |
| **k<sub>3</sub>** | LP and MT | 0.009 | 0.010 | 0.914 | 42 | 0.517 | -0.011 | 0.028 |
|  | LP and PC | 0.000 | 0.010 | 0.041 | 42 | 0.967 | -0.019 | 0.020 |
|  | MT and PC | -0.008 | 0.010 | -0.872 | 42 | 0.517 | -0.028 | 0.011 |
| **k<sub>4</sub>** | LP and MT | -0.014 | 0.014 | -0.965 | 42 | 0.517 | -0.043 | 0.015 |
|  | LP and PC | -0.008 | 0.014 | -0.527 | 42 | 0.687 | -0.037 | 0.022 |
|  | MT and PC | 0.006 | 0.014 | 0.438 | 42 | 0.724 | -0.023 | 0.036 |
| **Delay** | LP and MT | -0.859 | 0.158 | -5.440 | 42 | <0.001* | -1.178 | -0.541 |
|  | LP and PC | -0.303 | 0.158 | -1.916 | 42 | 0.166 | -0.621 | 0.016 |
|  | MT and PC | 0.557 | 0.158 | 3.524 | 42 | 0.003* | 0.238 | 0.875 |
| **V<sub>T</sub>** | LP and MT | 0.048 | 0.051 | 0.953 | 42 | 0.517 | -0.054 | 0.151 |
|  | LP and PC | -0.030 | 0.051 | -0.586 | 42 | 0.687 | -0.132 | 0.073 |
|  | MT and PC | -0.078 | 0.051 | -1.540 | 42 | 0.305 | -0.181 | 0.024 |
| **DVR** | LP and MT | 0.023 | 0.023 | 0.960 | 42 | 0.517 | -0.025 | 0.070 |
|  | LP and PC | -0.013 | 0.023 | -0.545 | 42 | 0.687 | -0.060 | 0.035 |
|  | MT and PC | -0.035 | 0.023 | -1.505 | 42 | 0.305 | -0.083 | 0.012 |

**Supplement Table 2:**
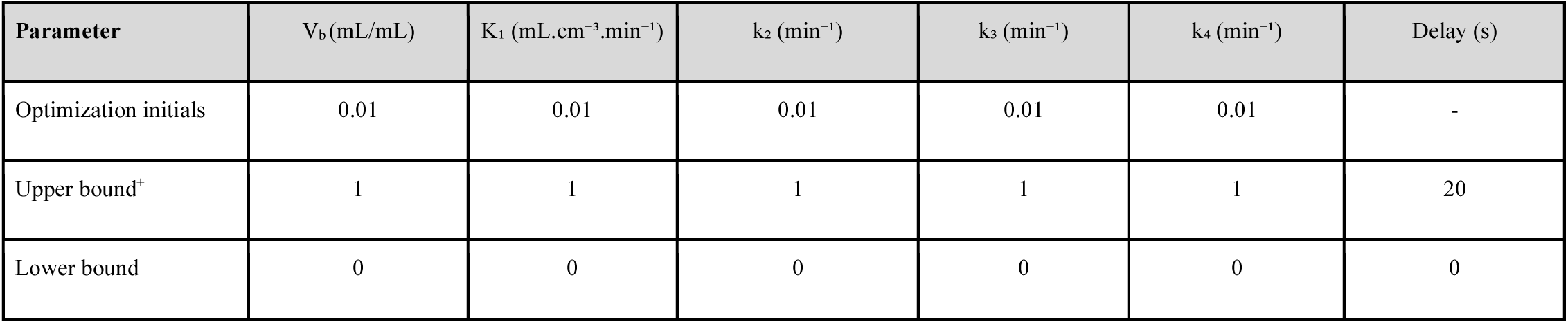
Description of optimization in our modeling. ^+^The upper bound 1 is chosen for all the parameters across all scan durations, except for the 10-minute dynamic duration. For the 10-minute scan duration, we observed instances where some parameter values exceeded 1, which is higher than the normal range observed in all subjects across all scan durations. Hence, we adjusted the upper bound for 10-minute scan duration as follows: V_b_ = 1, K_1_ = 5, k_2_ = 5, k_3_ = 5, and k_4_ = 5. The lower bound and optimization initials were as mentioned in the table.

**Supplement Figure 1:**
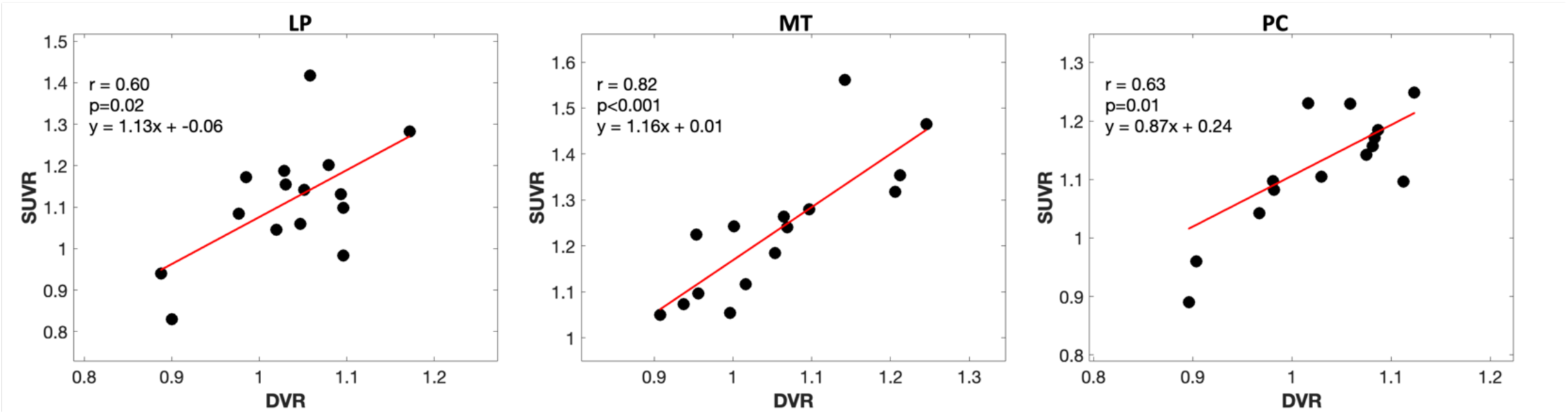
Association between SUVR (from 60 to 90 min) and DVR measures across ROIs.

**Supplement Figure 2:**
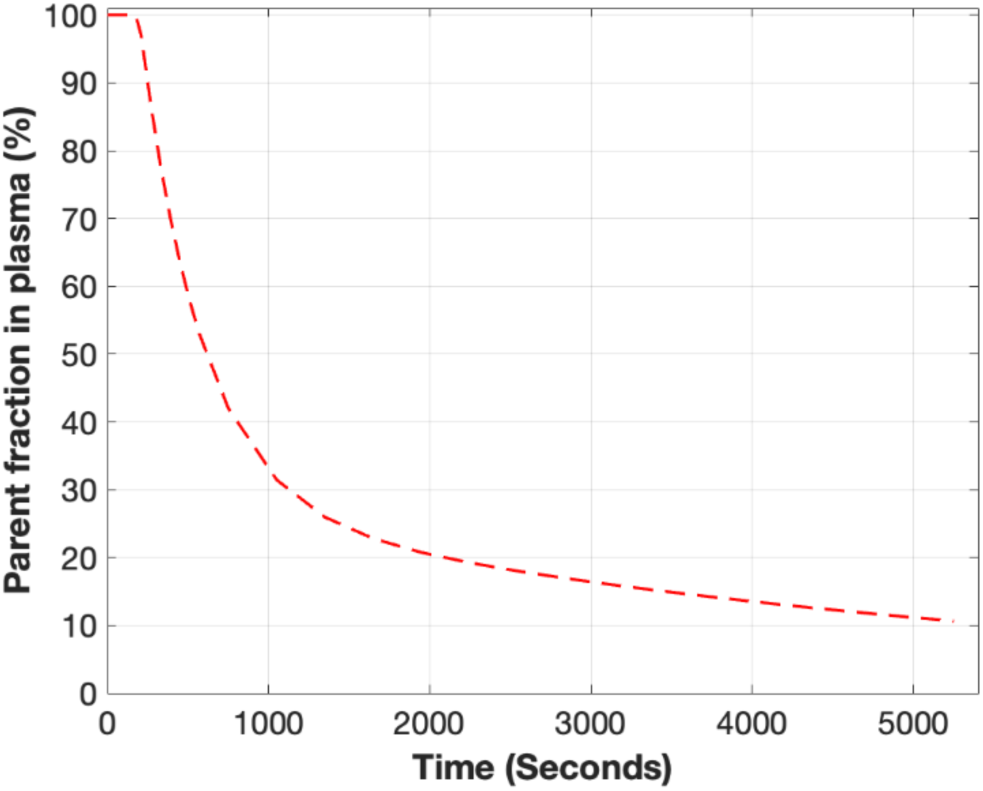
Illustrative example of the parent fraction in venous plasma over time was described with a biexponential function with the following parameters: α=72.01, τ_1_=6.38 min, τ_2_=87.15 min, t_0_ = 3.23 min, as discussed by Mueller et al.,2020 ^4^.

## Notes

### Competing Interest Statement

The authors have declared no competing interest.

## References

1. Kroth H, Oden F, Molette J, et al. Discovery and preclinical characterization of [18F]PI-2620, a next-generation tau PET tracer for the assessment of tau pathology in Alzheimer’s disease and other tauopathies. Eur J Nucl Med Mol Imaging 2019; 46: 2178–2189.

2. Song M, Beyer L, Kaiser L, et al. Binding characteristics of [18F]PI-2620 distinguish the clinically predicted tau isoform in different tauopathies by PET. J Cereb Blood Flow Metab 2021; 41: 2957–2972.

3. Bullich S, Barret O, Constantinescu C, et al. Evaluation of Dosimetry, Quantitative Methods, and Test-Retest Variability of 18F-PI-2620 PET for the Assessment of Tau Deposits in the Human Brain. J Nucl Med Off Publ Soc Nucl Med 2020; 61: 920–927.

4. Mueller A, Bullich S, Barret O, et al. Tau PET imaging with 18F-PI-2620 in Patients with Alzheimer Disease and Healthy Controls: A First-in-Humans Study. J Nucl Med Off Publ Soc Nucl Med 2020; 61: 911–919.

5. Wu M, Chen Z, Jiang M, et al. Friend or foe: role of pathological tau in neuronal death. Mol Psychiatry 2023; 28: 2215–2227.

6. Mormino EC, Toueg TN, Azevedo C, et al. Tau PET imaging with 18F-PI-2620 in aging and neurodegenerative diseases. Eur J Nucl Med Mol Imaging 2021; 48: 2233–2244.

7. Bischof GN, Brendel M, Barthel H, et al. Improved Tau PET SUVR Quantification in 4-Repeat Tau Phenotypes with [ ^18^ F]PI-2620. J Nucl Med 2024; 65: 952–955.

8. Bullich S, Mueller A, De Santi S, et al. Evaluation of tau deposition using 18F-PI-2620 PET in MCI and early AD subjects—a MissionAD tau sub-study. Alzheimers Res Ther 2022; 14: 105.

9. van der Weijden CWJ, Mossel P, Bartels AL, et al. Non-invasive kinetic modelling approaches for quantitative analysis of brain PET studies. Eur J Nucl Med Mol Imaging 2023; 50: 1636– 1650.

10. Volpi T, Maccioni L, Colpo M, et al. An update on the use of image-derived input functions for human PET studies: new hopes or old illusions? EJNMMI Res 2023; 13: 97.

11. Zanotti-Fregonara P, Chen K, Liow J-S, et al. Image-derived input function for brain PET studies: many challenges and few opportunities. J Cereb Blood Flow Metab 2011; 31: 1986– 1998.

12. Wang Y, Li E, Cherry SR, et al. Total-Body PET Kinetic Modeling and Potential Opportunities Using Deep Learning. PET Clin 2021; 16: 613–625.

13. Badawi RD, Shi H, Hu P, et al. First Human Imaging Studies with the EXPLORER Total-Body PET Scanner*. J Nucl Med 2019; 60: 299–303.

14. Spencer BA, Berg E, Schmall JP, et al. Performance Evaluation of the uEXPLORER Total-Body PET/CT Scanner Based on NEMA NU 2-2018 with Additional Tests to Characterize PET Scanners with a Long Axial Field of View. J Nucl Med Off Publ Soc Nucl Med 2021; 62: 861–870.

15. Holy EN, Li E, Bhattarai A, et al. Non-invasive quantification of 18F-florbetaben with total-body EXPLORER PET. EJNMMI Res 2024; 14: 39.

16. Song M, Scheifele M, Barthel H, et al. Feasibility of short imaging protocols for [18F]PI-2620 tau-PET in progressive supranuclear palsy. Eur J Nucl Med Mol Imaging 2021; 48: 3872– 3885.

17. Braak H, Braak E. Neuropathological stageing of Alzheimer-related changes. Acta Neuropathol (Berl) 1991; 82: 239–259.

18. Nardo L, Abdelhafez YG, Spencer BA, et al. Clinical Implementation of Total-Body PET/CT at University of California, Davis. PET Clin 2021; 16: 1–7.

19. Jenkinson M, Bannister P, Brady M, et al. Improved optimization for the robust and accurate linear registration and motion correction of brain images. NeuroImage 2002; 17: 825–841.

20. Fischl B. FreeSurfer. NeuroImage 2012; 62: 774–781.

21. Fischl B, Salat DH, Busa E, et al. Whole Brain Segmentation: Automated Labeling of Neuroanatomical Structures in the Human Brain. Neuron 2002; 33: 341–355.

22. Desikan RS, Ségonne F, Fischl B, et al. An automated labeling system for subdividing the human cerebral cortex on MRI scans into gyral based regions of interest. NeuroImage 2006; 31: 968–980.

23. Bhattarai A, Holy EN, Li E, et al. Comparison of delay correction methods to estimate 18F-florbetaben kinetics in the brain using an image derived input function on total-body EXPLORER PET. Alzheimers Dement 2023; 19: e079502.

24. Wang G, Nardo L, Parikh M, et al. Total-Body PET Multiparametric Imaging of Cancer Using a Voxelwise Strategy of Compartmental Modeling. J Nucl Med Off Publ Soc Nucl Med 2022; 63: 1274–1281.

25. Innis RB, Cunningham VJ, Delforge J, et al. Consensus Nomenclature for in vivo Imaging of Reversibly Binding Radioligands. J Cereb Blood Flow Metab 2007; 27: 1533–1539.

26. Logan J. Graphical analysis of PET data applied to reversible and irreversible tracers. Nucl Med Biol 2000; 27: 661–670.

27. Young CB, Winer JR, Younes K, et al. Divergent Cortical Tau Positron Emission Tomography Patterns Among Patients With Preclinical Alzheimer Disease. JAMA Neurol 2022; 79: 592– 603.

28. Lowe VJ, Bruinsma TJ, Wiste HJ, et al. Cross-sectional associations of tau-PET signal with cognition in cognitively unimpaired adults. Neurology; 93. Epub ahead of print 2 July 2019. DOI: 10.1212/WNL.0000000000007728.

29. Morris ED, Endres CJ, Schmidt KC, et al. Kinetic Modeling in Positron Emission Tomography. In: Emission Tomography. Elsevier, pp. 499–540.

30. Meindl M, Zatcepin A, Gnörich J, et al. Assessment of [18F]PI-2620 Tau-PET Quantification via Non-Invasive Automatized Image Derived Input Function. Eur J Nucl Med Mol Imaging. Epub ahead of print 8 May 2024. DOI: 10.1007/s00259-024-06741-7.

31. Providência L, Van Der Weijden CWJ, Mohr P, et al. Can Internal Carotid Arteries Be Used for Noninvasive Quantification of Brain PET Studies? J Nucl Med 2024; 65: 600–606.

32. Sari H, Mingels C, Alberts I, et al. First results on kinetic modelling and parametric imaging of dynamic 18F-FDG datasets from a long axial FOV PET scanner in oncological patients. Eur J Nucl Med Mol Imaging 2022; 49: 1997–2009.

33. Wierenga CE, Hays CC, Zlatar ZZ. Cerebral Blood Flow Measured by Arterial Spin Labeling MRI as a Preclinical Marker of Alzheimer’s Disease. J Alzheimers Dis JAD 2014; 42: S411– S419.

34. Bangen KJ, Thomas KR, Sanchez DL, et al. Entorhinal Perfusion Predicts Future Memory Decline, Neurodegeneration, and White Matter Hyperintensity Progression in Older Adults. J Alzheimers Dis JAD 2021; 81: 1711–1725.

